# Ecological indicators predict functional diversity dynamics following glacier retreat

**DOI:** 10.1101/2025.03.20.644423

**Authors:** Gianalberto Losapio, Lucía Mottet, Nora Khelidj, Bao Ngan Tu, Bruno E.L. Cerabolini, Stephanie Grand, Natasha de Vere, Antoine Guisan

**Affiliations:** Institute of Earth Surface Dynamics, University of Lausanne, 1015 Lausanne, Switzerland; Department of Biosciences, University of Milan, 20133 Milan, Italy; Department of Biotechnologies and Life Sciences, University of Insubria, Varese, Italy; Natural History Museum of Denmark, University of Copenhagen, Copenhagen, Denmark; Department of Ecology and Evolution, University of Lausanne, Lausanne, Switzerland

**Author notes:** **Corresponding author:** Gianalberto Losapio, University of Milan.

**Keywords:** alpine ecosystems, biodiversity change, climate change, community dynamics, glacier retreat, global warming, Landolt values, plant functional traits

## Abstract

The retreat of glaciers due to climate change is reshaping mountain landscapes and biodiversity. While previous research has documented vegetation succession after glacier retreat, our understanding of functional diversity dynamics is still limited. In this case study, we address the effects of glacier retreat on plant functional diversity by integrating plant traits with ecological indicator values across a 140-year chronosequence in a subalpine glacier landscape. We reveal that functional richness and functional dispersion decrease with glacier retreat, while functional evenness and functional divergence increase, suggesting a shift towards more specialized and competitive communities. Our findings highlight the critical role of ecological factors related to soil moisture, soil nutrients, and light availability in shaping plant community dynamics. The integrative approach of this case study provides novel insights into the potential response of alpine plant communities to climate change, offering a deeper understanding of how to predict and anticipate the effects of glacier extinction on biodiversity in rapidly changing environments.

## Introduction

Increasing global temperatures are causing the current retreat of glaciers worldwide (IPCC, 2022, 2019). Glacier retreat has been increasingly documented as a key driver of ecological change in mountain ecosystems, affecting the evolution of populations and the structure of communities and landscapes (Milner et al., 2017, 2007). Glacier retreat also leads to the emergence of new habitats characterized by barren, nutrient-poor terrains that are gradually colonized by diverse plant species that replace one another over space and time (Bosson et al., 2023; Cantera et al., 2024; Chapin et al., 1994; Ficetola et al., 2021). This process of primary succession not only alters species composition but also drives changes in plant functional traits (Caccianiga et al., 2006; Khelidj et al., 2024; Losapio et al., 2021), which are critical for both species persistence and ecosystem functioning (Brown et al., 2018; Cadotte, 2017; Losapio and Schöb, 2017). While the general patterns of plant primary succession following glacier retreat are relatively well understood, the mechanisms by which functional traits respond to the newly created environments remain less explored. Understanding how functional diversity responds to glacier retreat and ecological changes is crucial for predicting the future of biodiversity, especially under scenarios of continued warming and loss of glaciers.

Functional traits are measurable properties of plants that influence their fitness and performance in specific environments (Díaz et al., 2016; Garnier et al., 2016; Pérez-Harguindeguy et al., 2013). Traits include morphological characteristics, such as leaf area and canopy height, and reflect physiological conditions like leaf nitrogen content and specific leaf area. Functional traits are increasingly recognized as powerful tools for understanding ecological processes because they directly link species’ phenotype to ecological functions, such as nutrient cycling, productivity, and resilience to environmental stressors (Ackerly, 2004; Bongers et al., 2021; Cadotte, 2017; Losapio and Schöb, 2017). In the context of glacier retreat, functional traits can provide insights into the response of plants to rapidly changing conditions (Caccianiga et al., 2006; Ricotta et al., 2016). For example, traits that confer drought tolerance such as high leaf dry mater content or enable efficient nutrient uptake like low species leaf area are likely to be advantageous in the nutrient-poor and moisture-limited environments of glacier forelands. Additionally, understanding how these traits vary in plant communities along environmental gradients (Guisan et al., 2019) and across different successional stages (Khelidj et al., 2024) can help anticipate which functional changes will affect communities in these changing environments over time. Despite the growing interest in functional traits, how functional diversity changes with glacier retreat over spacetime remain poorly quantified, limiting our understanding of the consequences of glacier retreat on ecological processes and ecosystem functions.

Functional diversity (FD), which encompasses the community-weighted mean, range, and distribution of functional traits are critical for assessing ecological functions and strategies of multiple traits, species, and communities at once (Cadotte et al., 2011; Gross et al., 2009; Ricotta et al., 2016), going beyond traditional taxonomic analyses. High FD often indicates a robust ecosystem with a wide range of functional strategies, potentially leading to greater stability and resilience in the face of environmental disturbances and stressors. Conversely, low FD can suggest a dominance of few functional strategies (Laliberté and Legendre, 2010; Mason et al., 2012, 2005), potentially making the ecosystem more vulnerable to changes in environmental conditions. When environmental conditions change rapidly and species are colonizing and replacing each other, like in the case of glacier forelands, functional diversity can provide important insights into the processes of community assembly and ecosystem development (Brown et al., 2018; Gross et al., 2009; Shipley et al., 2017).

A multi-faceted approach to FD analysis provides specific ecological insights, from the diversity of available functional roles (functional richness) to the uniformity of resource use (functional evenness), degree of niche differentiation (functional divergence), and the distribution of functional strategies (functional dispersion) (Mason et al., 2005; Schleuter et al., 2010). Such integrated approach allows to investigate both the extent and distribution of functional trait space occupied by plant communities after glacier retreat (Khelidj et al., 2024). However, the dynamics of FD following glacier retreat remain poorly understood and difficult to predict, particularly in relation to changes of specific ecological conditions that follow glacier retreat and ecological succession.

Yet, it is important to recognize that community development and vegetation succession is shaped by other latent environmental factors beyond time alone, including soil and climate conditions (Burga et al., 2010; Charles et al., 2024; Ficetola et al., 2021; Losapio et al., 2021; Raffl et al., 2006; Tu et al., 2024; Walker et al., 2010). A possible solution to the challenge of considering changes in soil and climate along with time is inferring latent ecological conditions by means of bioindication principles (Diekmann, 2003). Ecological indicator values (EIVs) (Landolt et al., 2010; Scherrer and Guisan, 2019) can help not only to quantify the environmental preferences of species, for example for humidity, temperature, and soil nutrient, but also and especially as bioindicators (Fan et al., 2025; Scherrer et al., 2024). This way, EIVs such as those developed by Landolt and colleagues for the Swiss flora (Landolt et al., 2010) provide a standardized method for both assessing how species respond to environmental gradients as well as to inferring latent environmental conditions (Diekmann, 2003; Scherrer and Guisan, 2019; Shipley et al., 2017). Combining EIVs with FD analysis allows gaining a more comprehensive understanding of the adaptive strategies and ecological functions of plant communities (Wenskus et al., 2025).

This case study provides a novel framework for understanding the processes of community assembly and how functional strategies shift in the rapidly changing environmental conditions created by the retreat of glaciers during the last century. We integrate EIV with plant traits across different stages of succession following the retreat of a subalpine glacier. We address the following research questions: (1) How do EIVs and FD change after glacier retreat? (2) How do EIVs predict FD along with time over the succession? Following the successional gradient patterns (Bazzaz et al., 1990; Caccianiga et al., 2006; Chapin et al., 1994) and the ‘peak biodiversity’ concept (Losapio et al., 2025), we hypothesize that intermediate stages of succession promote species with fast-growth strategies, which are gradually replaced by species with traits enhancing competitive ability and resource conservation towards the end of succession ; Khelidj et al., 2024; Losapio et al., 2021). This case study aims to contribute not only to our understanding of primary succession after glacier retreat but also to provide novel insights into understanding the broader implications of climate change for mountain biodiversity and ecological processes.

## Methods

### Study site

The Mont Miné glacier foreland in Val d’Hérens, Switzerland (46°3’33.646’N, 7°32’54.550’E) was selected for this case study given its well preserved moraine system (Lambiel et al., 2016; Nicolussi et al., 2022) and its minimal elevation variation (1961–2000 m a.s.l.). The Mont Miné glacier has retreated by approximately 2.53 km since the end of the Little Ice Age (*c* 1860) as of 2023 (GLAMOS, 2023). Given its topography and hence the absence of an elevation gradient that covaries with terrain age, this glacier foreland offers a valuable system for investigating the impacts of climate change on biodiversity and ecological communities.

Integrating geochronological information on glacier dynamics from the catchment (Lambiel et al., 2016; Nicolussi et al., 2022) with historical cartographic reconstructions (Federal Office of Topography, 2023; GLAMOS, 2023) and field validation, we reconstructed a chronosequence to capture different stages of ecosystem development through time and across the glacier foreland by estimating terrain age corresponding to the time that has passed since the glacier retreated. Four major chronosequence stages were identified along the glacier foreland, which represent terrains exposed since at least the year 1989, 1925, 1900, and 1864. The average age of each stage was calculated as the mean between two adjacent moraines, resulting in the following terrain ages: 17 years for Stage 1, 66 years for Stage 2, 111 years for Stage 3, and 141 years for Stage 4. Sampling was conducted exclusively on the west side of the foreland to avoid disturbances from hydropower activities on the eastern side (Lambiel et al., 2016).

### Data collection

Within each stage, we randomly selected four sampling sites, totaling 16 sites. At each site, four 5 m × 5 m quadrats were established, covering an area of 100 m² per stage. Within each quadrat, plant communities were surveyed by recording species composition and estimating plant cover with a visual accuracy of 10%. Species identification followed *Flora Alpina* (Aeschimann et al., 2004). In total, we recorded 127 plant species across 32 families (Supplementary Table 10).

As ecological indicators, Landolt EIVs were considered as they are the most accurate and complete in describing soil factors and climate factors for the examined flora (Landolt et al., 2010). The following Landolt EIVs for soil factors were considered: (i) Soil moisture (F, here called H) ranging from very dry (1) to water-saturated soils (5); (ii) Soil reaction (R) ranging from extreme acidity with pH 3–4.5 (1) to basic reaction with pH > 6.5 (5); (iii) Soil nutrient (N) ranging from extremely infertile (1) to over-fertilized soils (5). The following Landolt EIVs for climate factors were considered: (i) Light availability (L) ranging from plants growing in shade with less than 3% relative illumination (1) to full sun (5); Air temperature (T), for plants typical of cold environment (1) to indicators of warmer places (5); Continentality (K, here called C) for plants whose center of distribution ranges from regions with oceanic climate (1) to continental climate (5). Although we do not expect macroscale differences in climate along the examined gradient, we considered climate-related EIVs like T and K to explore potential changes in microclimate conditions that might be reflected by the local plant community. After extracting Landolt EIVs and cross-referencing with the results of the plant community survey, 93 plant species with complete EIVs were retained for the statistical analysis (Supplementary Table 11).

We considered a set of ten plant functional traits representing key ecological strategies relevant for plant growth, reproduction, and nutrient cycling (Díaz et al., 2016; Garnier et al., 2016; Gross et al., 2009; Shipley et al., 2017; Wright et al., 2005). The following traits were considered: lateral spread (LS), canopy height (CH), leaf nitrogen content (LNC), leaf carbon content (LCC), leaf carbon-to-nitrogen ratio (CN), leaf area (LA), specific leaf area (SLA), leaf fresh weight (LFW), leaf dry weight (LDW), and leaf dry-matter content (LDMC). These leaf traits were obtained by integrating trait database (Kattge et al., 2020) with public, published data (Dalle Fratte et al., 2021; Khelidj et al., 2024; Losapio et al., 2021). After cross-referencing these datasets with the plant community survey, 53 plant species out of 93 with complete trait and EIVs data were finally retained for the statistical analysis (Supplementary Table 12). For each trait, we calculated the mean value for each species across all specimens to produce species-level trait means.

### Functional diversity

To assess changes in FD across successional stages in the Mont Miné glacier foreland and in relation to EIVs, we selected FD indices that reflect different aspects of community structure: functional richness (FRic), functional evenness (FEve), functional divergence (FDiv), and functional dispersion (FDis). Each index captures a different facet of FD (Mason et al., 2005; Schleuter et al., 2010) including dimensions of richness (the extent of niche space occupied), evenness (distribution within that space), and divergence (degree of species differentiation). Their combined use provides both diverse insights into and a comprehensive picture of how different functional strategies coexist over time and contribute to ecosystem processes following glacier retreat. These indices were calculated based on species-level trait means using the *FD* R package (Laliberté et al., 2014; Laliberté and Legendre, 2010).

FRic represents the total volume of functional trait space occupied by the community, and is equivalent to the convex hull volume of the species in trait space (Villéger et al., 2008). Ecologically, FRic reflects the range of functional roles within a community and indicates the diversity of strategies available for resource use, competition, and adaptation. Higher FRic values suggest a broader niche space, potentially enhancing ecosystem resilience by supporting a variety of ecological functions (Cornwell and Ackerly, 2009; Mason et al., 2005).

FEve measures the evenness of trait distribution within the functional space, and was calculated using a minimum spanning tree which connects all species based on their trait distances (Villéger et al., 2008). The higher the FEve the more regular the distribution of species along resource gradients, suggesting efficient and balanced resource use across the community. In contrast, low FEve values imply clustering in specific areas of trait space, potentially indicating competitive dominance or environmental filtering (Mason et al., 2012)

FDiv quantifies the degree of divergence from the centroid in trait space, focusing on species that deviate most from average community traits. This index highlights niche differentiation, where higher FDiv values indicate greater specialization and ecological complementarity among species (Villéger et al., 2008). FDiv is particularly useful for identifying communities where species have distinct functional roles as it underscores the degree to which species occupy different ecological niches to minimize niche overlap (Mason et al., 2005).

FDis represents the mean distance of each species from the centroid in trait space, weighted by relative abundance (Laliberté and Legendre, 2010). Unlike other indices, FDis is independent of species richness, making it an effective metric for comparing communities with varying species numbers. High FDis values suggest a wide distribution of functional strategies, potentially contributing to ecosystem resilience and adaptability. FDis is valuable for capturing the functional breadth of a community and understanding how species abundances relate to trait variability (Laliberté & Legendre, 2010).

### Data analysis

To investigate the ecological conditions and functional composition of plant communities, we calculated community-weighted means (CWMs) for each EIV and functional trait. CWMs represent the mean value of an EIV or a functional trait within a community, weighted by the relative abundance of each species. This approach provides quantitative insights into community-level conditions and strategies, and their responses to glacier retreat and ecological gradients. CWMs were calculated as

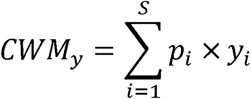

where *p_i_* is the relative abundance of each species *i* occurring in a community composed of *S* species, and *y_i_* is the EIV or functional trait value for the species *i*. This analysis was performed for each response variable *y* (i.e., a single EIV or functional trait) across all sampling plots and successional stages, allowing us to assess how EIV and FD shift in relation to glacier retreat. CWMs were calculated using the *FD* package in R (Laliberté et al., 2014; Laliberté and Legendre, 2010), ensuring consistency with other FD analyses.

To address the first research question, we used linear regression models to test for differences in each *CWM_y_* and FD index (response variables in separate models) across the succession (predictor, categorical variable with Stage 1 as reference level) (Fox and Weisser, 2019). Type-II Analysis of Variance was applied to the fitted model to calculate F-tests for predictor significance using the *car* R package (Fox and Weisser, 2019). Inference of model parameters was conducted using the Wald method, which computes confidence intervals and associated *p*-values by dividing the parameter estimate (intercept and slope) by its standard error, then comparing this statistic against a *t-*distribution with residual degrees of freedom (Lüdecke et al., 2020). Cohen’s *f* was calculated as effect size to assess the magnitude of change in each index across stages (Ben-Shachar et al., 2020), helping to interpret the ecological relevance of observed trends across indices.

To address the second research question and assess the multivariate relationships among glacier retreat, EIVs, functional traits, and FD, we first conducted a Principal Component Analysis (PCA) using the *FactoMineR* package in R (Lê et al., 2008). PCA was chosen to reduce data dimensionality while preserving the primary variation in the dataset, enabling us to visualize and interpret the key ecological gradients and functional trait variation. After calculating the proportion of variance explained by each PC, we identified the variables that were significantly associated with as well as contributing the most to the first two PC by looking at correlation coefficients between variables and PC using the *dimdesc* function of *FactoMineR* (Lê et al., 2008). In a preliminary, exploratory way, we also considered the strength and direction of univariate relationships between EIVs and FD (Supplementary Fig. S2; Supplementary Table 8).

Then, to make inference on the ecological drivers of FD, we used two complementary approaches: a parametric model (linear regression) and a non-parametric, machine learning algorithm (random forest). In both cases, CWM of functional traits and functional diversity indices were considered as response variables with respect to glacier retreat (years since glacier retreat) while CWM_s_ of EIVs (H, R, N, L, T, C), which were used as predictors.

An initial linear model was fitted for each response variable using all predictor variables. To refine each model and identify the most significant predictors, we performed stepwise model selection based on Akaike Information Criterion (AIC) using the *stepAIC* function of *MASS* R package (Venables and Ripley, 2002). This approach systematically removed non-significant predictors, yielding a more parsimonious model. Although this procedure is highly sensitive to the order of the predictors in the model, given the relatively low number of predictors and their non-collinearity, stepwise selection would still provide an efficient approach for this case study. For each final model, we evaluated the effects of the selected predictors using type-II analysis of variance (F-test), inferring model parameters with 95% CIs and *p*-values, and calculating Cohen’s *f* standardized effect sizes.

Finally, to further examine the relative importance of EIVs for predicting FD, we implemented a Random Forest (RF) model with permutation testing to assess the statistical significance of variable importance. The Random Forest model was chosen for its ability to handle complex, non-linear relationships and for being robust to assumptions on data distribution. As mentioned above, functional traits and FD indices were modeled as a function of years since glacier retreat and EIVs. The *rfPermute* R package (Archer, 2023) was used to build the model, which included 5,000 decision trees on 500 bootstrap samples of the initial data to provide robust significance testing for variable importance, yielding 2’500’000 samples. Following model training, we summarized the results to evaluate model performance (see Appendix R script). For each model, the importance and significance of predictors was assessed based on the percentage of increase in mean square error when their values are randomly permuted (Δ*MSE*). To summarize predictor importance over all FD indices, we calculated the average predictor importance index 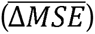.

## Results

Four out of the six ecological indicator values (EIVs) significantly changed with successional stages (Supplementary Table 1). Cohen’s effect size of glacier retreat was large for soil moisture (*f* = 1.26), soil reaction (*f* = 1.93), soil nutrient (*f* = 1.27), and light availability (*f =* 1.93); it was medium but unsignificant for air temperature (*f* = 0.49) and continentality (*f* = 0.81).

Looking at the direction of changes and trends (Fig. 1; Supplementary Table 2), CWMs of EIVs for soil moisture increase with glacier retreat at the latest, 140-years successional stage (β = 0.38 [0.17 – 0.59], *p =* 0.002). On the contrary, CWM_s_ of EIVs for soil reaction decrease with glacier retreat at the 110-years stage (β = - 0.70 [ - 1.13 – - 0.27], *p =* 0.004) and 140-years stage (β = - 1.22 [− 1.65 – - 0.79], *p <* 0.001). CWM_s_ of EIVs for soil nutrient marginally decrease at the 65-years stage (β = - 0.25 [− 0.51 – 0.01], *p =* 0.061) and 110- years stage (β = - 0.25 [ - 0.52 – 0.01], *p =* 0.060), while significantly decrease with glacier retreat at the 140-years stage (β =- 0.53 [ - 0.80 – - 0.27], *p =* 0.001). CWM_s_ of EIVs for light availability marginally decrease at 110-years stage (β = - 0.45 [− 0.90 – 0.01], *p =* 0.052) and significantly decrease with glacier retreat at 140-years stage (β =- 1.20 [− 1.65 – - 0.75], *p <* 0.001). CWMs of EIVs for air temperature show no evidence of significant directional change associated to glacier retreat stages. CWMs of EIVs for continentality decrease with glacier retreat at the latest, 140-years successional stage (β = −0.32 [− 0.59 – - 0.59], *p =* 0.002).

**Fig. 1.**
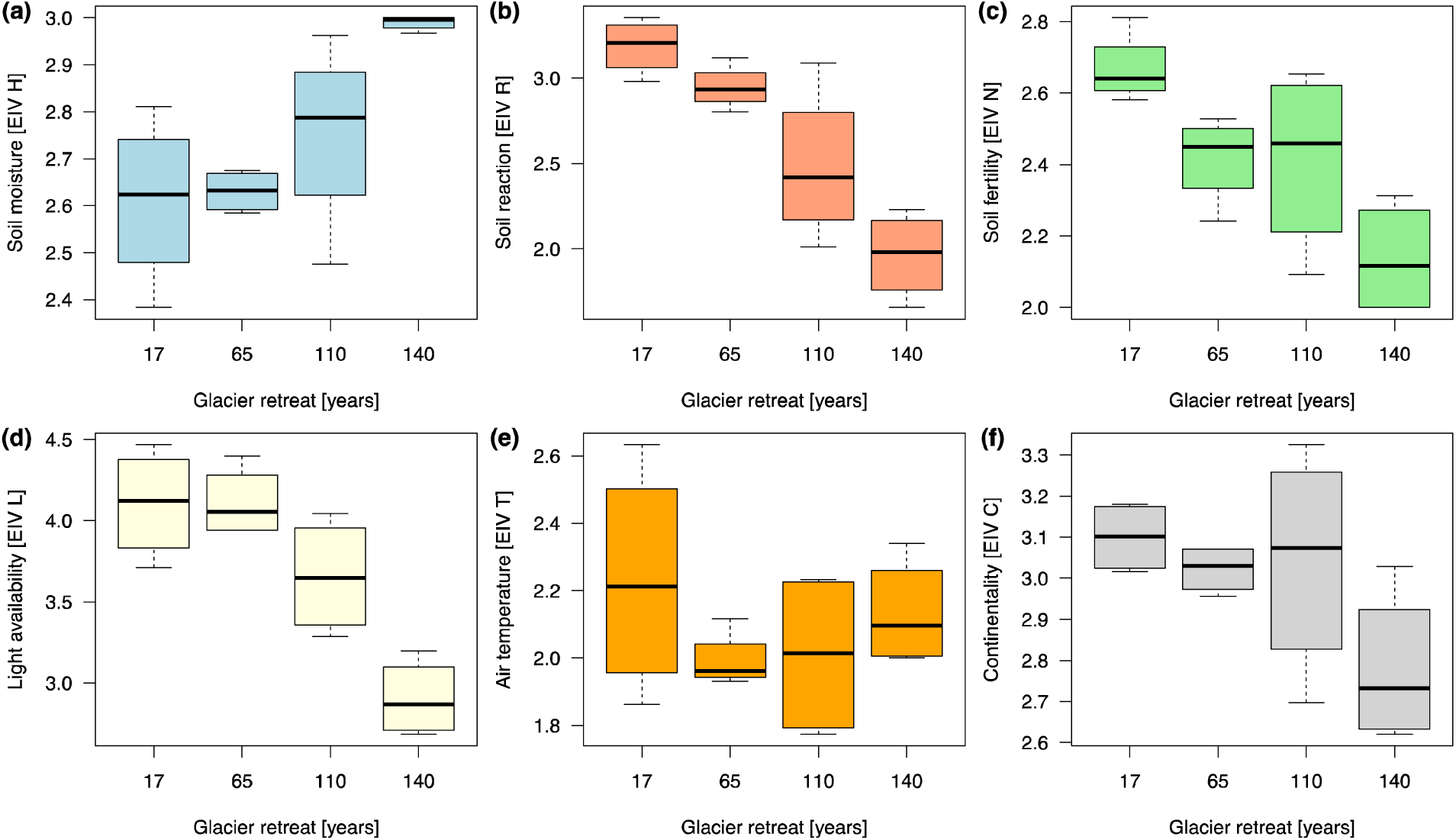
Community-Weighted Means of Landolt Ecological Indicator Values (EIVs) of plant communities along the ecological succession triggered by glacier retreat.

Nine out of ten plant functional traits significantly or marginally (one) changed along the succession (Supplementary Table 3). Cohen’s effect size of glacier retreat was large for lateral spread (*f* = 1.11), leaf nitrogen content (LNC, *f* = 0.93), leaf carbon content (LCC, *f* = 2.17), leaf carbon:nitrogen ration (CN, *f* = 1.18), leaf area (LA, *f* = 1.00), leaf dry weight (LDW, *f* = 1.32), leaf fresh weight (LFW, *f* = 1.27), leaf dry matter content (LDMC, *f* = 1.36), and canopy height (*f* = 1.74); it was medium but unsignificant for specific leaf area (SLA, *f* = 0.64).

Looking at the direction of change and trends of plant functional traits (Fig. 2; Supplementary Table 2), CWMs of lateral spread, canopy height, LCC, CN, LDW, and LDMC significantly increase with glacier retreat with a minimum detectable change at the latest 140-years stage. On the contrary, LA, LFW, and SLA were the highest at the intermediate stages of 65-years and 110-years (Supplementary Table 2).

**Fig. 2.**
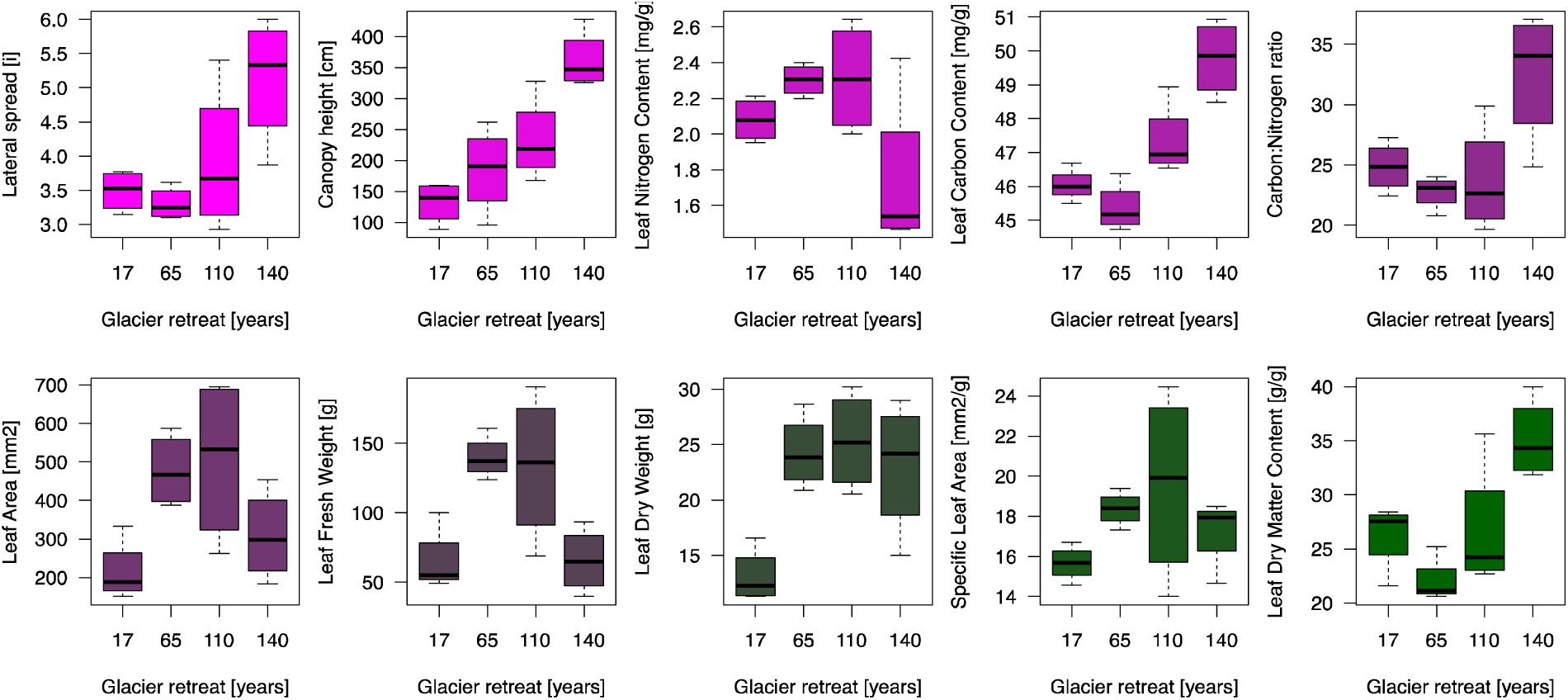
Plant functional traits CWM along the ecological succession triggered by glacier retreat.

One FD index out of four, namely FDis, was significantly associated to succession overall (Supplementary Table 5). Cohen’s effect size of glacier retreat was large for FDis (*f* = 1.37), and it was medium but unsignificant for FRic (*f* = 1.83), FEve (*f* = 0.60), and FDiv (*f* = 0.57). Yet, both FRic (Fig. 3a) and FDis (Fig. 3d) significantly decrease with glacier retreat at the latest, 140-years successional stage (β = - 5.32 [− 10.10 – 0.54], *p =* 0.032, and β = - 0.44 [− 0.67 – - 0.21], *p =* 0.001, respectively). Furthermore, both FEve (Fig. 3b) and FDiv (Fig. 3c) marginally increase at the 140-years successional stage (β = 0.10 [− 0.01 – 0.21], *p =* 0.062, and β = 0.12 [− 0.02 – 0.26], *p =* 0.091, respectively).

**Fig. 3.**
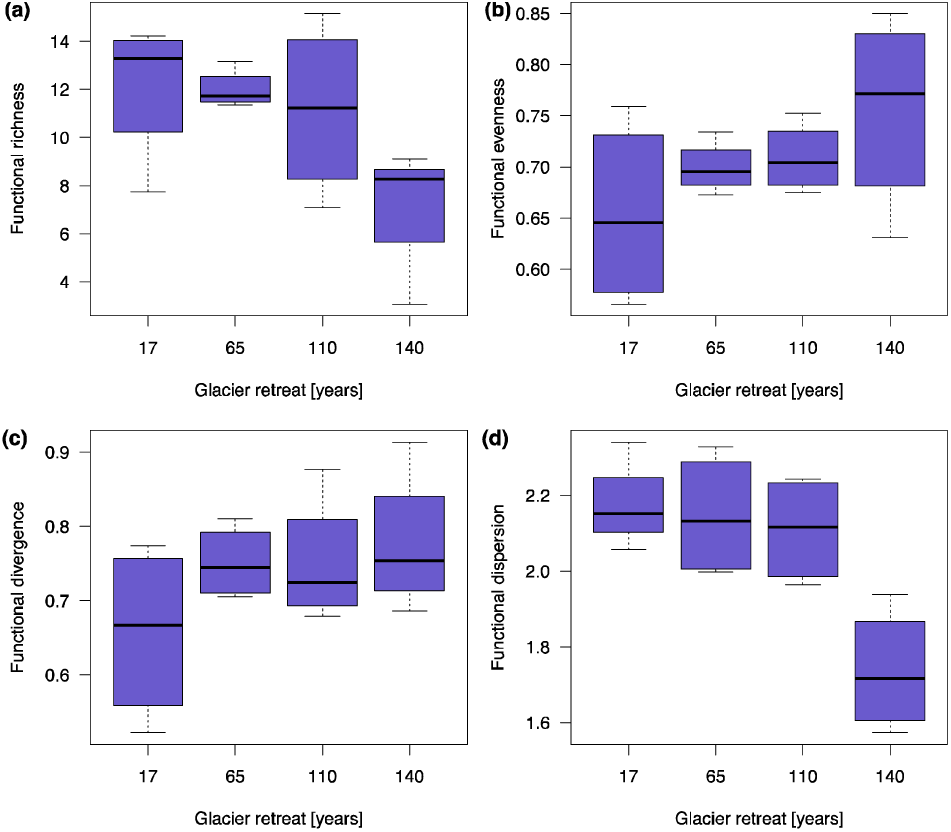
Functional diversity indices along the ecological succession triggered by glacier retreat.

To address the second question, the Principal Component Analysis (Fig. 4; Supplementary Table 7) identified major patterns in the variation of EIVs and FD as glacier retreats. The first component (PC1) and the second component (PC2) accounted for 50.8% and 20.3% of the variance, respectively, hence explaining together 71.1% of the total variance. Key variables positively contributing to PC1 are functional traits LCC, LS, LDMC, CAN, CN as well as EIVs of soil moisture and years since glacier retreat; EIVs of soil reaction, soil nutrient, light availability, and continentality as well as LNC, LFW, FDis and FRic are negatively associated with PC1. PC2 is positively associated with traits including LDW, LA, LFW, SLA, LNC as well as years since glacier retreat and FDiv. These trends reflect a major gradient in soil development which captures differences in resource allocation, carbon economy and plant size, separating pioneer and late plant communities adapted to high versus low soil nutrient and light environments, respectively (Supplementary Fig. S1).

**Fig. 4.**
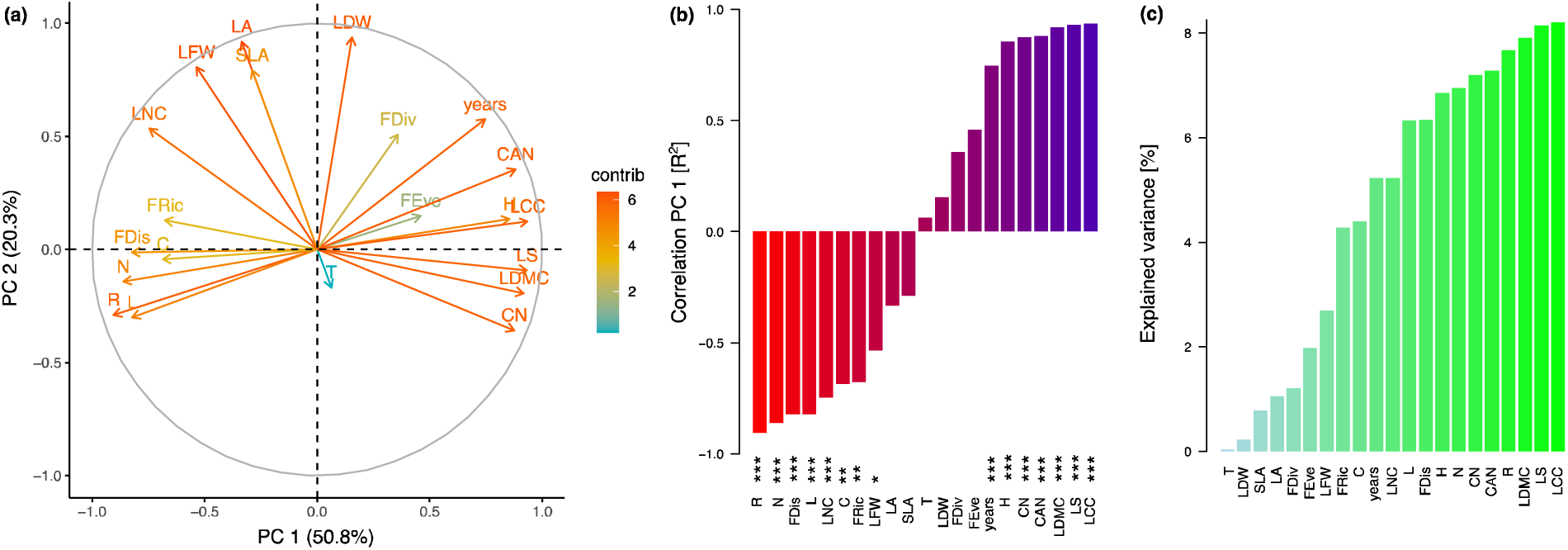
Principal component analysis of the relationships among glacier retreat (years), ecological indicators (soil humidity ‘H’, soil ph ‘R’, soil nutrients ‘N’, light ‘L’, temperature ‘T’, continentality ‘C’), plant traits (Lateral Spread ‘LS’, Leaf Nitrogen Content ‘LNC’, Leaf Carbon Content ‘LCC’, leaf Carbon:Nitrogen ratio ‘CN’, Canopy height ‘CAN’), and FD indices (Functional Dispersion ‘FDis’, Functional Richness ‘FRic’, Function Evenness ‘FEve’, Functional Divergence ‘FDiv’). (a) Factor map of the variables contributing to the first two components. The color gradient represents the contribution of variables. Map of the sites is reported in Supplementary Fig. S1. (b) Correlation of the variables with the first component. Significance of *p*-values is reported as following: 0.05<*p<*0.01=*, 0.01<*p<*0.001=**, *p<*0.001=*** (c) Contributions of variables according to their explained variance.

To address the third question, the final model for functional traits (n = 10) included glacier retreat (years) eight times out of ten, soil nutrient EIVs seven times, light availability EIVs six times, soil moisture EIVs twice, EIVs of soil reaction and continentality once, while air temperature EIVs was never selected as major predictor in any final model (Fig. 5a; Supplementary Table 8). For functional diversity indices (n = 4), the final model included glacier retreat (years), soil nutrient EIVs, and light availability EIVs two times out of four, soil moisture EIVs, soil reaction EIVs, and continentality once, and air temperature EIVs none (Fig. 5a; Supplementary Table 8). Overall, on average, soil nutrient EIVs had the strongest effects on FD (*f̅* = 0.52), followed by glacier retreat (*f̅* = 0.39), light availability EIVs (*f̅* = 0.29), and soil moisture EIVS (*f̅* = 0.14).

**Fig. 5.**
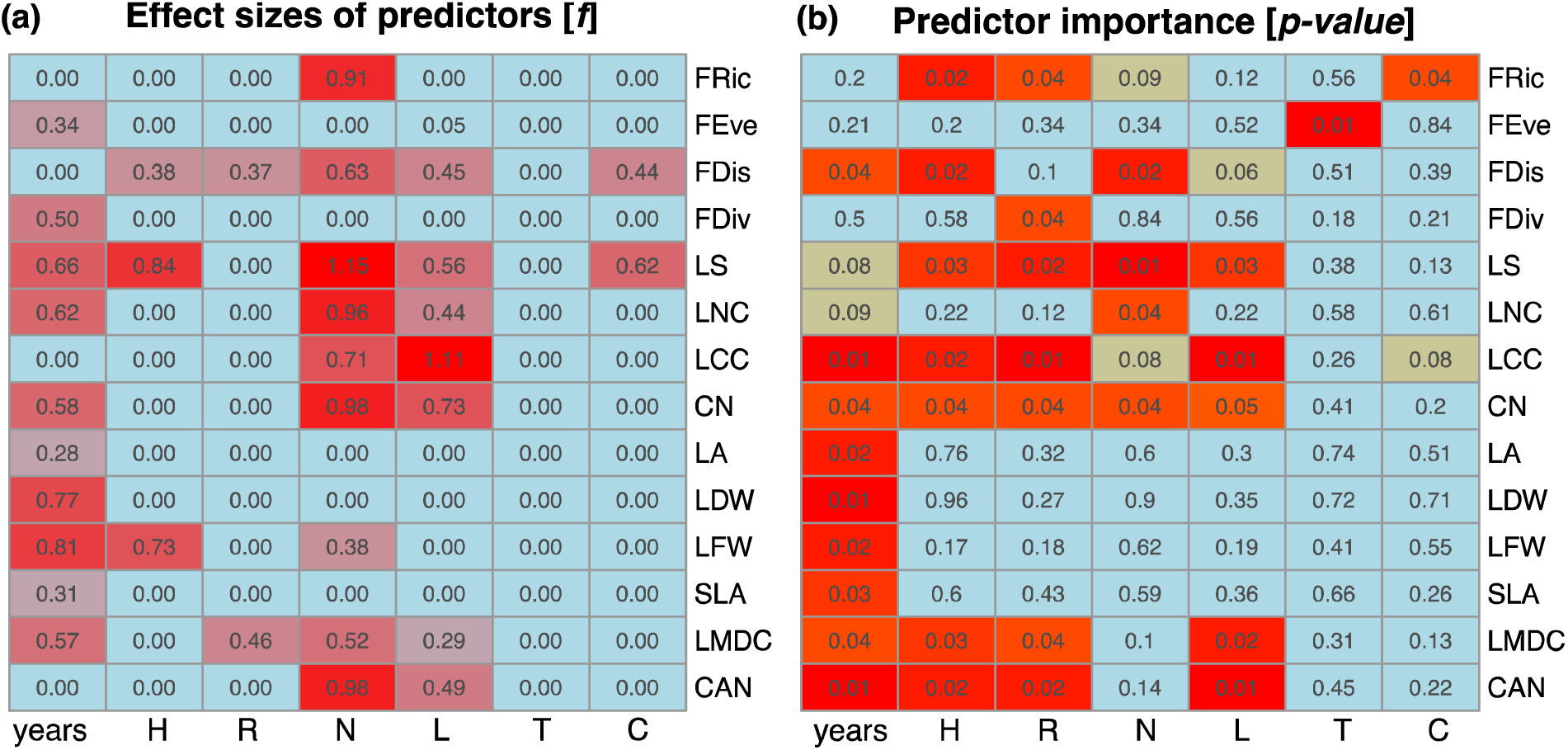
Ecological indicator values (soil humidity ‘H’, soil ph ‘R’, soil nutrients ‘N’, light ‘L’, temperature ‘T’, continentality ‘C’) and glacier retreat (years) as predictors of plant traits (Lateral Spread ‘LS’, Leaf Nitrogen Content ‘LNC’, Leaf Carbon Content ‘LCC’, leaf Carbon:Nitrogen ratio ‘CN’, Canopy height ‘CAN’), and FD indices (Functional Dispersion ‘FDis’, Functional Richness ‘FRic’, Function Evenness ‘FEve’, Functional Divergence ‘FDiv’). (a) Cohen’s *f* effect size estimating how much variance in FD variables is accounted for by EIVs in the final regression model. This model, built upon stepwise selection according to AIC, includes major predictors as highlighted in reddish colors and excludes less significant predictors, which are highlighted in light blue. (b) Variable importance in the Random Forest model measured via the percentage of increase in mean square errors, proportional to color gradient intensity; scores indicate the associated *p-*values.

Qualitatively similar results were obtained using the random forest algorithm (Fig. 5b; Supplementary Table 9). Glacier retreat (years) was among the most important predictors for eight functional traits and one functional diversity index (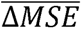 = 17.2%); soil moisture and soil reaction EIVs were among the most important predictors for five functional traits and two functional diversity indices (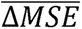 = 13.2% and 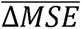 = 16.5%, respectively); light availability EIVs was among the most important predictors for five functional traits but no functional diversity index (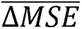 = 14.1%); soil nutrient EIVs was among the most important predictors for three functional traits and one functional diversity index (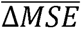 = 10.6%); air temperature and continentality EIVs were among the most important predictors for one functional diversity index and no functional traits (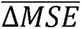 = 0.5% and 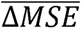 = 4.7%, respectively.

Finally, it is worth noticing that both FRic and FDis increase with increasing soil nutrient EIVs (β = 9.58 [3.28 – 15.89], *p =* 0.006; and (β = 0.80 [− 0.09 – 1.69], *p =* 0.074). Soil nutrient EIVs is also positively associated with increasing LNC (β = 1.31 [0.45 – 2.18], *p =* 0.006) while negatively associated with LS (β =- 2.71 [ - 4.37 – - 1.05], *p =* 0.005), LCC (β =- 3.02 [ - 5.58 – - 0.47], *p =* 0.024), CN (β =- 16.78 [ - 27.54 – - 6.01], *p =* 0.005), and CAN (β =- 2.54 [ - 4.08 – - 0.99], *p =* 0.004).

## Discussion

Our study sheds new light on the relationships between glacier retreat, ecological indicator values (EIVs), and plant functional diversity within a subalpine glacier ecosystem, providing novel insights into the factors driving community assembly in response to rapidly changing environmental conditions. We demonstrate that glacier retreat drives substantial shifts FD, with EIVs playing a crucial role in predicting these changes. Particularly, soil-related EIVs including soil nutrient availability, moisture, and pH exhibited significant directional shifts over successional stages, reflecting changes in resource availability and habitat conditions. FD metrics responded in distinct ways: functional richness (FRic) and functional dispersion (FDis) declined with succession, suggesting a reduction in the range of available functional strategies over time, while functional evenness (FEve) and functional divergence (FDiv) showed marginal increases, indicating greater niche differentiation in later stages. Additionally, regression and machine learning models highlighted the dominant role of soil nutrient and light availability EIVs in shaping FD, while climatic indicators such as air temperature and continentality were weak predictors of functional trait variation. These findings suggest that microhabitat conditions and soil development exert a stronger influence on community assembly and dynamics than macroclimatic factors, emphasizing the importance of integrating EIVs with FD to better understand vegetation responses to glacier retreat and climate change.

These findings reveal that functional diversity evolves predictably after glacier retreat, highlighting the critical role of ecological factors, particularly soil nutrient and light availability, in shaping plant community dynamics and functional trait distributions over space and time. These results contribute to a deeper understanding of the functional strategies of plant communities in response to climate change and glacier retreat. Indicators such as soil moisture, soil reaction, soil nutrient, and light availability were highly responsive to glacier retreat, as evidenced by large effect sizes, indicating significant shifts in ecological conditions over time. Soil moisture indicators increased at later successional stages, suggesting that progressive soil development enhances water retention capacities via pioneer herbaceous species modification of bedrock. This increase in soil moisture conditions suggests that early stages are comparatively drier despite the higher water availability provided by glacier melting and alluvial sediments. In contrast, indicators of soil nutrient, reaction and light availability decreased, likely due to the accumulation of organic matter, the litter deposition and canopy shading by late-successional shrubs and coniferous trees, which host few plant species adapted to acidic, low-light environments.

These shifts in EIVs align with previous studies documenting successional changes in soil properties and vegetation in glacier forelands (Bernasconi et al., 2011; Burga et al., 2010; Charles et al., 2024), where initial colonizers gradually create conditions favorable for more competitive species (Caccianiga et al., 2006; Ficetola et al., 2021; Khelidj et al., 2024; Losapio et al., 2021). Yet, we report a clear decrease in soil nutrient EIVs despite the potential increase in soil nutrient content (Charles et al., 2024), a pattern that may be due to rapid nutrient uptake and nitrogen sequestration by competitive species (Caccianiga et al., 2006). This process is further evidenced below by shifts in functional traits related to carbon and nitrogen economy in late stages. The distinct trajectories of EIVs provide a nuanced perspective on how soil development and resource availability act as selective pressures for community assembly and dynamics throughout succession in this study.

Nine out of ten plant functional traits showed significant or marginal changes following glacier retreat, reflecting changes in species composition and inherent ecological responses to the changing environmental conditions. Traits such as lateral spread, canopy height, leaf carbon content, leaf carbon:nitrogen content and LDMC increased with succession, suggesting a shift towards species with larger size and greater carbon allocation and nutrient sequestration, which are advantageous in more stable, resource-enriched environments. In contrast, traits associated with rapid resource acquisition, such as leaf area, SLA, and leaf nitrogen content were highest at intermediate successional stages. These findings align with the hypothesis that intermediate stages of succession promote species with fast-growth strategies, which are gradually replaced by species with traits enhancing competitive ability and resource conservation towards the end of succession (Bazzaz et al., 1990; Caccianiga et al., 2006; Chapin et al., 1994; Khelidj et al., 2024; Losapio et al., 2021). Indeed, these trait dynamics highlight a functional shift from resource-acquisitive to resource-conservative strategies, a trend documented in successional studies within glacier forelands and other primary successions (Burga et al., 2010; Cutler et al., 2008; Fastie, 2013; Ficetola et al., 2021).

Our findings indicate that functional dispersion (FDis) was the strongest index associated to succession, as evidenced by a large effect size and a marked decrease at the latest stage of 140 years. The decline in FDis suggests that the range of trait combinations within the community narrows with glacier retreat, which could reflect a shift towards a more specialized set of functional strategies (Laliberté and Legendre, 2010; Mason et al., 2012). This trend suggests increased niche partitioning and competitive exclusion as the habitat stabilizes (e.g., lower physical/topographic disturbance) and becomes more productive (e.g., higher soil fertility). While functional richness (FRic) also decreased significantly at the 140-year stage, its overall association with succession was weaker, with a medium but non-significant effect size. This pattern suggests that, although the total volume of functional space occupied by species initially expands as communities diversify, it ultimately contracts in later stages as fewer species dominate. Interestingly, functional evenness (FEve) and functional divergence (FDiv) showed marginal increases in the later successional stages, suggesting a shift towards greater resource use complementarity and niche differentiation (Mason et al., 2005; Villéger et al., 2008). This pattern is consistent with previous studies suggesting that functional diversity declines as competitive exclusion intensifies in more stable stages (Cadotte, 2017; Garnier et al., 2016; Khelidj et al., 2024).

The decrease in functional richness and dispersion, coupled with an increase in evenness and divergence, underscores the potential dual role of functional diversity in early successional stages: promoting resilience by accommodating a range of functional strategies while also enabling long-term stability through niche partitioning. These trends support the hypothesis that functional diversity mediates ecosystem resilience by fostering a balance between redundancy and specialization within plant communities (Bongers et al., 2021; Díaz et al., 2016).

Notably, EIVs could predict functional traits and functional diversity, indicating that variation in FD was strongly influenced by shifting ecological conditions, particularly soil nutrient, soil moisture, soil reaction, and light availability. These indicators thus suggest that ecological factors associated to soil and canopy development act as critical ecological mechanisms shaping and selecting both the composition and functionality of plant communities over space and time. These results reinforce the notion that FD is closely tied to environmental gradients (Diekmann, 2003; Guisan et al., 2019; Losapio and Schöb, 2017; Scherrer and Guisan, 2019), suggesting that the development of ecosystems is driven by trait selection to the specific ecological conditions created by glacier retreat. The stronger predicting power of EIVs for soil conditions and light along with the weak association of air temperature and continentality EIVs with functional diversity might reflect the buffering effect of microhabitats, and the importance of local biotic factors in shaping community assembly and dynamics.

Interesting enough, EIVs of large-scale climate factors such as those associated to air temperature and continentality fall short in predicting FD. This result is not surprising given the absence of covariation between terrain age and elevation, which may potentially confound the effects of glacier retreat by climate on its own, and given the low climatic variability over the Mont Minè glacier foreland. The results of this case study suggests that while these climatic variables undoubtedly influence broader ecological patterns, their direct impact on the specific functional strategies of plant communities in the examined glacier foreland may be overshadowed by more local environmental factors such as soil pH, light conditions, and nutrient availability.

By linking FD with EIVs, we provide a novel perspective on how plant communities assemble, adapt, and function under rapidly changing environmental conditions. Yet, future research should incorporate wider, long-term monitoring and additional environmental variables, such as soil microbiota and microclimate variation. It should also extend to other altitudinal and biogeographical ranges to fully understand the adaptive potential of mountain ecosystems facing glacier extinction.

In conclusion, the integration of EIVs with FD offers a comprehensive approach to understanding ecological factors underlying community assembly and adaptation while forecasting broader ecological changes. The results of this case study have broader implications for the future of biodiversity in mountain ecosystems considering increasing rates of glacier retreat. The ecological dynamics we observed here may extend to other mountain landscapes, highlighting the importance of monitoring FD together with ecological indicators while offering valuable insights into understanding how plant communities respond and potentially adapt to ongoing climate change.

## Author Contributions

GL conceived and designed the study. LM and GL contributed to data generation and analysis. NK, BNT, BELC, NDV, SG, and AG contributed with designing the study, data collection, field expertise, and manuscript drafting. All authors contributed to the interpretation of results. GL drafted the manuscript and all authors read and approved the final version.

## Acknowledgments

We thank the Service des forêts, de la nature et du paysage of Canton du Valais for providing sampling permission. We thank the Biodiversity Change group for providing feedbacks.

## Appendix

Supplementary information with summary tables.

## Data Availability

Data and R script (no novel code) will be published on Dataverse and GitHub, respectively, upon manuscript acceptance.

## References

Ackerly, D., 2004. Functional strategies of chaparral shrubs in relation to seasonal water deficit and disturbance. Ecol. Monogr. 74, 25–44. 10.1890/03-4022

Aeschimann, D., Lauber, K., Moser, D.M., Theurillat, J.-P., 2004. Flora alpina. Haupt, Bern.

Archer, E., 2023. rfPermute: Estimate Permutation p-Values for Random Forest Importance Metrics.

Bazzaz, F.A., Grace, J.B., Tilman, D., 1990. 12 - Plant–Plant Interactions in Successional Environments, in: Perspectives on Plant Competition. Academic Press, pp. 239–263. 10.1016/B978-0-12-294452-9.50016-3

Ben-Shachar, M.S., Lüdecke, D., Makowski, D., 2020. effectsize: Estimation of Effect Size Indices and Standardized Parameters. J. Open Source Softw. 5, 2815. 10.21105/joss.02815

Bernasconi, S.M., Bauder, A., Bourdon, B., Brunner, I., Bünemann, E., Chris, I., Derungs, N., Edwards, P., Farinotti, D., Frey, B., Frossard, E., Furrer, G., Gierga, M., Göransson, H., Gülland, K., Hagedorn, F., Hajdas, I., Hindshaw, R., Ivy-Ochs, S., Jansa, J., Jonas, T., Kiczka, M., Kretzschmar, R., Lemarchand, E., Luster, J., Magnusson, J., a.D. Mitchell, E., Venterink, H.O., Plötze, M., Reynolds, B., Smittenberg, R.H., Stähli, M., Tamburini, F., Tipper, E.T., Wacker, L., Welc, M., Wiederhold, J.G., Zeyer, J., Zimmermann, S., Zumsteg, A., 2011. Chemical and Biological Gradients along the Damma Glacier Soil Chronosequence, Switzerland. Vadose Zone J. 10, 867. 10.2136/vzj2010.0129

Bongers, F.J., Schmid, B., Bruelheide, H., Bongers, F., Li, S., von Oheimb, G., Li, Y., Cheng, A., Ma, K., Liu, X., 2021. Functional diversity effects on productivity increase with age in a forest biodiversity experiment. Nat. Ecol. Evol. 5, 1594–1603. 10.1038/s41559-021-01564-3

Bosson, J.B., Huss, M., Cauvy-Fraunié, S., Clément, J.C., Costes, G., Fischer, M., Poulenard, J., Arthaud, F., 2023. Future emergence of new ecosystems caused by glacial retreat. Nature 620, 562–569. 10.1038/s41586-023-06302-2

Brown, L.E., Khamis, K., Wilkes, M., Blaen, P., Brittain, J.E., Carrivick, J.L., Fell, S., Friberg, N., Füreder, L., Gislason, G.M., Hainie, S., Hannah, D.M., James, W.H.M., Lencioni, V., Olafsson, J.S., Robinson, C.T., Saltveit, S.J., Thompson, C., Milner, A.M., 2018. Functional diversity and community assembly of river invertebrates show globally consistent responses to decreasing glacier cover. Nat. Ecol. Evol. 2, 325–333. 10.1038/s41559-017-0426-x

Burga, C.A., Krüsi, B., Egli, M., Wernli, M., Elsener, S., Ziefle, M., Fischer, T., Mavris, C., 2010. Plant succession and soil development on the foreland of the Morteratsch glacier (Pontresina, Switzerland): Straight forward or chaotic? Flora - Morphol. Distrib. Funct. Ecol. Plants 205, 561–576.

Caccianiga, M., Luzzaro, A., Pierce, S., Ceriani, R.M., Cerabolini, B., 2006. The functional basis of a primary succession resolved by CSR classification. Oikos 112, 10–20. 10.1111/j.0030-1299.2006.14107.x

Cadotte, M.W., 2017. Functional traits explain ecosystem function through opposing mechanisms. Ecol. Lett. 20, 989–996. 10.1111/ele.12796

Cadotte, M.W., Carscadden, K., Mirotchnick, N., 2011. Beyond species: Functional diversity and the maintenance of ecological processes and services. J. Appl. Ecol. 48, 1079–1087. 10.1111/j.1365-2664.2011.02048.x

Cantera, I., Carteron, A., Guerrieri, A., Marta, S., Bonin, A., Ambrosini, R., Anthelme, F., Azzoni, R.S., Almond, P., Alviz Gazitúa, P., Cauvy-Fraunié, S., Ceballos Lievano, J.L., Chand, P., Chand Sharma, M., Clague, J., Cochachín Rapre, J.A., Compostella, C., Cruz Encarnación, R., Dangles, O., Eger, A., Erokhin, S., Franzetti, A., Gielly, L., Gili, F., Gobbi, M., Hågvar, S., Khedim, N., Meneses, R.I., Peyre, G., Pittino, F., Rabatel, A., Urseitova, N., Yang, Y., Zaginaev, V., Zerboni, A., Zimmer, A., Taberlet, P., Diolaiuti, G.A., Poulenard, J., Thuiller, W., Caccianiga, M., Ficetola, G.F., 2024. The importance of species addition ‘versus’ replacement varies over succession in plant communities after glacier retreat. Nat. Plants 10, 256–267. 10.1038/s41477-023-01609-4

Chapin, F.S., Walker, L.R., Fastie, C.L., Sharman, L.C., 1994. Mechanisms of Primary Succession Following Deglaciation at Glacier Bay, Alaska. Ecol. Monogr. 64, 149–175. 10.2307/2937039

Charles, C., Khelidj, N., Mottet, L., Tu, B.N., Adatte, T., Bomou, B., Faria, M., Monbaron, L., Reubi, O., de Vere, N., Grand, S., Losapio, G., 2024. Plant–Soil interactions drive development of novel ecosystems after glacier retreat. Res. Sq. 10.21203/rs.3.rs-4647918/v1

Cornwell, W.K., Ackerly, D.D., 2009. Community assembly and shifts in plant trait distributions across an environmental gradient in coastal California. Ecol. Monogr. 79, 109–126. 10.1890/07-1134.1

Cutler, N. a., Belyea, L.R., Dugmore, a. J., 2008. The spatiotemporal dynamics of a primary succession. J. Ecol. 96, 231–246. 10.1111/j.1365-2745.2007.01344.x

Dalle Fratte, M., Pierce, S., Zanzottera, M., Cerabolini, B.E.L., 2021. The association of leaf sulfur content with the leaf economics spectrum and plant adaptive strategies. Funct. Plant Biol. 48, 924–935.

Díaz, S., Kattge, J., Cornelissen, J.H.C., Wright, I.J., Lavorel, S., Dray, S., Reu, B., Kleyer, M., Wirth, C., Colin Prentice, I., Garnier, E., Bönisch, G., Westoby, M., Poorter, H., Reich, P.B., Moles, A.T., Dickie, J., Gillison, A.N., Zanne, A.E., Chave, J., Joseph Wright, S., Sheremet’ev, S.N., Jactel, H., Baraloto, C., Cerabolini, B., Pierce, S., Shipley, B., Kirkup, D., Casanoves, F., Joswig, J.S., Günther, A., Falczuk, V., Rüger, N., Mahecha, M.D., Gorné, L.D., 2016. The global spectrum of plant form and function. Nature 529, 167–171.

Diekmann, M., 2003. Species indicator values as an important tool in applied plant ecology – a review. Basic Appl. Ecol. 4, 493–506. 10.1078/1439-1791-00185

Fan, Y., Zhang, C., Hu, W., Khan, K.S., Zhao, Y., Huang, B., 2025. Development of soil quality assessment framework: A comprehensive review of indicators, functions, and approaches. Ecol. Indic. 172, 113272. 10.1016/j.ecolind.2025.113272

Fastie, C.L., 2013. Causes and Ecosystem Consequences of Multiple Pathways of Primary Succession at Glacier CAUSES AND ECOSYSTEM CONSEQUENCES OF MULTIPLE PATHWAYS OF PRIMARY SUCCESSION AT GLACIER BAY, ALASKA ‘.

Federal Office of Topography, 2023. Swisstopo.

Ficetola, G.F., Marta, S., Guerrieri, A., Gobbi, M., Ambrosini, R., Fontaneto, D., Zerboni, A., Poulenard, J., Caccianiga, M., Thuiller, W., 2021. Dynamics of Ecological Communities Following Current Retreat of Glaciers. Annu. Rev. Ecol. Evol. Syst. 10.1146/annurev-ecolsys-010521-040017

Fox, J., Weisser, S., 2019. An R companion to Applied Regression, 3rd ed. Sage, Thousand Oaks, CA.

Garnier, E., Navas, M.-L., Grigulis, K., 2016. Plant Functional Diversity. Oxford University Press.

GLAMOS, 2023. Swiss Glacier Volume Change. Glacier Monitoring Switzerland.

Gross, N., Kunstler, G., Liancourt, P., De Bello, F., Suding, K.N., Lavorel, S., 2009. Linking individual response to biotic interactions with community structure: A trait-based framework. Funct. Ecol. 23, 1167–1178. 10.1111/j.1365-2435.2009.01591.x

Guisan, A., Mod, H.K., Scherrer, D., Münkemüller, T., Pottier, J., Alexander, J.M., D’Amen, M., 2019. Scaling the linkage between environmental niches and functional traits for improved spatial predictions of biological communities. Glob. Ecol. Biogeogr. 28, 1384–1392. 10.1111/geb.12967

IPCC, 2022. Climate Change 2022: Impacts, Adaptation, and Vulnerability. Cambridge University Press.

IPCC, 2019. IPCC Special Report on the Ocean and Cryosphere in a Changing Climate. Cambridge University Press, Cambridge, UK and New York, NY, USA.

Kattge, J., Bönisch, G., Díaz, S., Lavorel, S., Prentice, I.C., Leadley, P., Tautenhahn, S., Werner, G.D.A., Aakala, T., Abedi, M., Acosta, A.T.R., Adamidis, G.C., Adamson, K., Aiba, M., Albert, C.H., Alcántara, J.M., Alcázar C, C., Aleixo, I., Ali, H., Amiaud, B., Ammer, C., Amoroso, M.M., Anand, M., Anderson, C., Anten, N., Antos, J., Apgaua, D.M.G., Ashman, T.-L., Asmara, D.H., Asner, G.P., Aspinwall, M., Atkin, O., Aubin, I., Baastrup-Spohr, L., Bahalkeh, K., Bahn, M., Baker, T., Baker, W.J., Bakker, J.P., Baldocchi, D., Baltzer, J., Banerjee, A., Baranger, A., Barlow, J., Barneche, D.R., Baruch, Z., Bastianelli, D., Battles, J., Bauerle, W., Bauters, M., Bazzato, E., Beckmann, M., Beeckman, H., Beierkuhnlein, C., Bekker, R., Belfry, G., Belluau, M., Beloiu, M., Benavides, R., Benomar, L., Berdugo-Lattke, M.L., Berenguer, E., Bergamin, R., Bergmann, J., Bergmann Carlucci, M., Berner, L., Bernhardt-Römermann, M., Bigler, C., Bjorkman, A.D., Blackman, C., Blanco, C., Blonder, B., Blumenthal, D., Bocanegra-González, K.T., Boeckx, P., Bohlman, S., Böhning-Gaese, K., Boisvert-Marsh, L., Bond, W., Bond-Lamberty, B., Boom, A., Boonman, C.C.F., Bordin, K., Boughton, E.H., Boukili, V., Bowman, D.M.J.S., Bravo, S., Brendel, M.R., Broadley, M.R., Brown, K.A., Bruelheide, H., Brumnich, F., Bruun, H.H., Bruy, D., Buchanan, S.W., Bucher, S.F., Buchmann, N., Buitenwerf, R., Bunker, D.E., Bürger, J., Burrascano, S., Burslem, D.F.R.P., Butterfield, B.J., Byun, C., Marques, M., Scalon, M.C., Caccianiga, M., Cadotte, M., Cailleret, M., Camac, J., Camarero, J.J., Campany, C., Campetella, G., Campos, J.A., Cano-Arboleda, L., Canullo, R., Carbognani, M., Carvalho, F., Casanoves, F., Castagneyrol, B., Catford, J.A., Cavender-Bares, J., Cerabolini, B.E.L., Cervellini, M., Chacón-Madrigal, E., Chapin, K., Chapin, F.S., Chelli, S., Chen, S.-C., Chen, A., Cherubini, P., Chianucci, F., Choat, B., Chung, K.-S., Chytrý, M., Ciccarelli, D., Coll, L., Collins, C.G., Conti, L., Coomes, D., Cornelissen, J.H.C., Cornwell, W.K., Corona, P., Coyea, M., Craine, J., Craven, D., Cromsigt, J.P.G.M., Csecserits, A., Cufar, K., Cuntz, M., da Silva, A.C., Dahlin, K.M., Dainese, M., Dalke, I., Dalle Fratte, M., Dang-Le, A.T., Danihelka, J., Dannoura, M., Dawson, S., de Beer, A.J., De Frutos, A., De Long, J.R., Dechant, B., Delagrange, S., Delpierre, N., Derroire, G., Dias, A.S., Diaz-Toribio, M.H., Dimitrakopoulos, P.G., Dobrowolski, M., Doktor, D., Dřevojan, P., Dong, N., Dransfield, J., Dressler, S., Duarte, L., Ducouret, E., Dullinger, S., Durka, W., Duursma, R., Dymova, O., E-Vojtkó, A., Eckstein, R.L., Ejtehadi, H., Elser, J., Emilio, T., Engemann, K., Erfanian, M.B., Erfmeier, A., Esquivel-Muelbert, A., Esser, G., Estiarte, M., Domingues, T.F., Fagan, W.F., Fagúndez, J., Falster, D.S., Fan, Y., Fang, J., Farris, E., Fazlioglu, F., Feng, Y., Fernandez-Mendez, F., Ferrara, C., Ferreira, J., Fidelis, A., Finegan, B., Firn, J., Flowers, T.J., Flynn, D.F.B., Fontana, V., Forey, E., Forgiarini, C., François, L., Frangipani, M., Frank, D., Frenette-Dussault, C., Freschet, G.T., Fry, E.L., Fyllas, N.M., Mazzochini, G.G., Gachet, S., Gallagher, R., Ganade, G., Ganga, F., García-Palacios, P., Gargaglione, V., Garnier, E., Garrido, J.L., de Gasper, A., Gea-Izquierdo, G., Gibson, D., Gillison, A.N., Giroldo, A., Glasenhardt, M.-C., Gleason, S., Gliesch, M., Goldberg, E., Göldel, B., Gonzalez-Akre, E., Gonzalez-Andujar, J.L., González-Melo, A., González-Robles, A., Graae, B.J., Granda, E., Graves, S., Green, W.A., Gregor, T., Gross, N., Guerin, G.R., Günther, A., Gutiérrez, A.G., Haddock, L., Haines, A., Hall, J., Hambuckers, A., Han, W., Harrison, S.P., Hattingh, W., Hawes, J.E., He, T., He, P., Heberling, J.M., Helm, A., Hempel, S., Hentschel, J., Hérault, B., Hereş, A.-M., Herz, K., Heuertz, M., Hickler, T., Hietz, P., Higuchi, P., Hipp, A.L., Hirons, A., Hock, M., Hogan, J.A., Holl, K., Honnay, O., Hornstein, D., Hou, E., Hough-Snee, N., Hovstad, K.A., Ichie, T., Igić, B., Illa, E., Isaac, M., Ishihara, M., Ivanov, L., Ivanova, L., Iversen, C.M., Izquierdo, J., Jackson, R.B., Jackson, B., Jactel, H., Jagodzinski, A.M., Jandt, U., Jansen, S., Jenkins, T., Jentsch, A., Jespersen, J.R.P., Jiang, G.-F., Johansen, J.L., Johnson, D., Jokela, E.J., Joly, C.A., Jordan, G.J., Joseph, G.S., Junaedi, D., Junker, R.R., Justes, E., Kabzems, R., Kane, J., Kaplan, Z., Kattenborn, T., Kavelenova, L., Kearsley, E., Kempel, A., Kenzo, T., Kerkhoff, A., Khalil, M.I., Kinlock, N.L., Kissling, W.D., Kitajima, K., Kitzberger, T., Kjøller, R., Klein, T., Kleyer, M., Klimešová, J., Klipel, J., Kloeppel, B., Klotz, S., Knops, J.M.H., Kohyama, T., Koike, F., Kollmann, J., Komac, B., Komatsu, K., König, C., Kraft, N.J.B., Kramer, K., Kreft, H., Kühn, I., Kumarathunge, D., Kuppler, J., Kurokawa, H., Kurosawa, Y., Kuyah, S., Laclau, J.-P., Lafleur, B., Lallai, E., Lamb, E., Lamprecht, A., Larkin, D.J., Laughlin, D., Le Bagousse-Pinguet, Y., le Maire, G., le Roux, P.C., le Roux, E., Lee, T., Lens, F., Lewis, S.L., Lhotsky, B., Li, Y., Li, X., Lichstein, J.W., Liebergesell, M., Lim, J.Y., Lin, Y.-S., Linares, J.C., Liu, C., Liu, D., Liu, U., Livingstone, S., Llusià, J., Lohbeck, M., López-García, Á., Lopez-Gonzalez, G., Lososová, Z., Louault, F., Lukács, B.A., Lukeš, P., Luo, Y., Lussu, M., Ma, S., Maciel Rabelo Pereira, C., Mack, M., Maire, V., Mäkelä, A., Mäkinen, H., Malhado, A.C.M., Mallik, A., Manning, P., Manzoni, S., Marchetti, Z., Marchino, L., Marcilio-Silva, V., Marcon, E., Marignani, M., Markesteijn, L., Martin, A., Martínez-Garza, C., Martínez-Vilalta, J., Mašková, T., Mason, K., Mason, N., Massad, T.J., Masse, J., Mayrose, I., McCarthy, J., McCormack, M.L., McCulloh, K., McFadden, I.R., McGill, B.J., McPartland, M.Y., Medeiros, J.S., Medlyn, B., Meerts, P., Mehrabi, Z., Meir, P., Melo, F.P.L., Mencuccini, M., Meredieu, C., Messier, J., Mészáros, I., Metsaranta, J., Michaletz, S.T., Michelaki, C., Migalina, S., Milla, R., Miller, J.E.D., Minden, V., Ming, R., Mokany, K., Moles, A.T., Molnár V, A., Molofsky, J., Molz, M., Montgomery, R.A., Monty, A., Moravcová, L., Moreno-Martínez, A., Moretti, M., Mori, A.S., Mori, S., Morris, D., Morrison, J., Mucina, L., Mueller, S., Muir, C.D., Müller, S.C., Munoz, F., Myers-Smith, I.H., Myster, R.W., Nagano, M., Naidu, S., Narayanan, A., Natesan, B., Negoita, L., Nelson, A.S., Neuschulz, E.L., Ni, J., Niedrist, G., Nieto, J., Niinemets, Ü., Nolan, R., Nottebrock, H., Nouvellon, Y., Novakovskiy, A., Network, T.N., Nystuen, K.O., O’Grady, A., O’Hara, K., O’Reilly-Nugent, A., Oakley, S., Oberhuber, W., Ohtsuka, T., Oliveira, R., Öllerer, K., Olson, M.E., Onipchenko, V., Onoda, Y., Onstein, R.E., Ordonez, J.C., Osada, N., Ostonen, I., Ottaviani, G., Otto, S., Overbeck, G.E., Ozinga, W.A., Pahl, A.T., Paine, C.E.T., Pakeman, R.J., Papageorgiou, A.C., Parfionova, E., Pärtel, M., Patacca, M., Paula, S., Paule, J., Pauli, H., Pausas, J.G., Peco, B., Penuelas, J., Perea, A., Peri, P.L., Petisco-Souza, A.C., Petraglia, A., Petritan, A.M., Phillips, O.L., Pierce, S., Pillar, V.D., Pisek, J., Pomogaybin, A., Poorter, H., Portsmuth, A., Poschlod, P., Potvin, C., Pounds, D., Powell, A.S., Power, S.A., Prinzing, A., Puglielli, G., Pyšek, P., Raevel, V., Rammig, A., Ransijn, J., Ray, C.A., Reich, P.B., Reichstein, M., Reid, D.E.B., Réjou-Méchain, M., de Dios, V.R., Ribeiro, S., Richardson, S., Riibak, K., Rillig, M.C., Riviera, F., Robert, E.M.R., Roberts, S., Robroek, B., Roddy, A., Rodrigues, A.V., Rogers, A., Rollinson, E., Rolo, V., Römermann, C., Ronzhina, D., Roscher, C., Rosell, J.A., Rosenfield, M.F., Rossi, C., Roy, D.B., Royer-Tardif, S., Rüger, N., Ruiz-Peinado, R., Rumpf, S.B., Rusch, G.M., Ryo, M., Sack, L., Saldaña, A., Salgado-Negret, B., Salguero-Gomez, R., Santa-Regina, I., Santacruz-García, A.C., Santos, J., Sardans, J., Schamp, B., Scherer-Lorenzen, M., Schleuning, M., Schmid, B., Schmidt, M., Schmitt, S., Schneider, J.V., Schowanek, S.D., Schrader, J., Schrodt, F., Schuldt, B., Schurr, F., Selaya Garvizu, G., Semchenko, M., Seymour, C., Sfair, J.C., Sharpe, J.M., Sheppard, C.S., Sheremetiev, S., Shiodera, S., Shipley, B., Shovon, T.A., Siebenkäs, A., Sierra, C., Silva, V., Silva, M., Sitzia, T., Sjöman, H., Slot, M., Smith, N.G., Sodhi, D., Soltis, P., Soltis, D., Somers, B., Sonnier, G., Sørensen, M.V., Sosinski Jr, E.E., Soudzilovskaia, N.A., Souza, A.F., Spasojevic, M., Sperandii, M.G., Stan, A.B., Stegen, J., Steinbauer, K., Stephan, J.G., Sterck, F., Stojanovic, D.B., Strydom, T., Suarez, M.L., Svenning, J.-C., Svitková, I., Svitok, M., Svoboda, M., Swaine, E., Swenson, N., Tabarelli, M., Takagi, K., Tappeiner, U., Tarifa, R., Tauugourdeau, S., Tavsanoglu, C., te Beest, M., Tedersoo, L., Thiffault, N., Thom, D., Thomas, E., Thompson, K., Thornton, P.E., Thuiller, W., Tichý, L., Tissue, D., Tjoelker, M.G., Tng, D.Y.P., Tobias, J., Török, P., Tarin, T., Torres-Ruiz, JoséM., Tóthmérész, B., Treurnicht, M., Trivellone, V., Trolliet, F., Trotsiuk, V., Tsakalos, J.L., Tsiripidis, I., Tysklind, N., Umehara, T., Usoltsev, V., Vadeboncoeur, M., Vaezi, J., Valladares, F., Vamosi, J., van Bodegom, P.M., van Breugel, M., Van Cleemput, E., van de Weg, M., van der Merwe, S., van der Plas, F., van der Sande, M.T., van Kleunen, M., Van Meerbeek, K., Vanderwel, M., Vanselow, K.A., V\aarhammar, A., Varone, L., Vasquez Valderrama, M.Y., Vassilev, K., Vellend, M., Veneklaas, E.J., Verbeeck, H., Verheyen, K., Vibrans, A., Vieira, I., Villacís, J., Violle, C., Vivek, P., Wagner, K., Waldram, M., Waldron, A., Walker, A.P., Waller, M., Walther, G., Wang, H., Wang, F., Wang, W., Watkins, H., Watkins, J., Weber, U., Weedon, J.T., Wei, L., Weigelt, P., Weiher, E., Wells, A.W., Wellstein, C., Wenk, E., Westoby, M., Westwood, A., White, P.J., Whitten, M., Williams, M., Winkler, D.E., Winter, K., Womack, C., Wright, I.J., Wright, S.J., Wright, J., Pinho, B.X., Ximenes, F., Yamada, T., Yamaji, K., Yanai, R., Yankov, N., Yguel, B., Zanini, K.J., Zanne, A.E., Zelený, D., Zhao, Y.-P., Zheng, Jingming, Zheng, Ji, Ziemińska, K., Zirbel, C.R., Zizka, G., Zo-Bi, I., Zotz, G., Wirth, C., 2020. TRY plant trait database –enhanced coverage and open access. Glob. Change Biol. 26, 119–188. 10.1111/gcb.14904

Khelidj, N., Caccianiga, M., Cerabolini, B.E.L., Tampucci, D., Losapio, G., 2024. Glacier extinction homogenizes functional diversity via ecological succession. J. Veg. Sci. 35, e13312. 10.1111/jvs.13312

Laliberté, E., Legendre, P., 2010. A distance-based framework for measuring functional diversity from multiple traits. Ecology 91, 299–305. 10.1890/08-2244.1

Laliberté, E., Legendre, P., Shipley, B., 2014. FD: Measuring Functional Diversity from Multiple Traits, and Other Tools for Functional Ecology. R Package Version 1.0-12.

Lambiel, C., Maillard, B., Kummert, M., Reynard, E., 2016. Geomorphology of the Hérens valley (Swiss Alps). J. Maps.

Landolt, E., Bäumler, B., Erhardt, A., Hegg, O., Klötzli, F., Lämmler, W., Nobis, M., Rudmann-Maurer, K., Schweingruber, F., Theurillat, J., et al, 2010. Flora indicativa. Ökologische Zeigerwerte und biologische Kennzeichen zur Flora der Schweiz und der Alpen. Ecological indicators values and biological attributes of the flora of Switzerland and the Alps, 2nd ed. Haupt, Bern, Switzerland.

Lê, S., Josse, J., Husson, F., 2008. FactoMineR: An R Package for Multivariate Analysis. J. Stat. Softw. 25, 1–18. 10.18637/jss.v025.i01

Losapio, G., Cerabolini, B.E.L., Maffioletti, C., Tampucci, D., Gobbi, M., Caccianiga, M., 2021. The Consequences of Glacier Retreat Are Uneven Between Plant Species. Front. Ecol. Evol. 8, 520. 10.3389/fevo.2020.616562

Losapio, G., Schöb, C., 2017. Resistance of plant–plant networks to biodiversity loss and secondary extinctions following simulated environmental changes. Funct. Ecol. 31, 1145–1152. 10.1111/1365-2435.12839

Lüdecke, D., Ben-Shachar, M.S., Patil, I., Makowski, D., 2020. Extracting, Computing and Exploring the Parameters of Statistical Models using R. J. Open Source Softw. 5, 2445. 10.21105/joss.02445

Mason, N.W.H., Mouillot, D., Lee, W.G., Wilson, J.B., 2005. Functional richness, functional evenness and functional divergence: the primary components of functional diversity. Oikos 111, 112–118. 10.1111/j.0030-1299.2005.13886.x

Mason, N.W.H., Richardson, S.J., Peltzer, D.A., de Bello, F., Wardle, D.A., Allen, R.B., 2012. Changes in coexistence mechanisms along a long-term soil chronosequence revealed by functional trait diversity. J. Ecol. 100, 678–689. 10.1111/j.1365-2745.2012.01965.x

Milner, A.M., Fastie, C.L., Chapin, F.S., Engstrom, D.R., Sharman, L.C., 2007. Interactions and Linkages among Ecosystems during Landscape Evolution. BioScience 57, 237–247. 10.1641/B570307

Milner, A.M., Khamis, K., Battin, T.J., Brittain, J.E., Barrand, N.E., Füreder, L., Cauvy-Fraunié, S., Gíslason, G.M., Jacobsen, D., Hannah, D.M., Hodson, A.J., Hood, E., Lencioni, V., Ólafsson, J.S., Robinson, C.T., Tranter, M., Brown, L.E., 2017. Glacier shrinkage driving global changes in downstream systems. Proc. Natl. Acad. Sci. 10.1073/pnas.1619807114

Nicolussi, K., Roy, M.L., Schlüchter, C., Stoffel, M., Wacker, L., 2022. The glacier advance at the onset of the Little Ice Age in the Alps: New evidence from Mont Miné and Morteratsch glaciers. The Holocene. 10.1177/09596836221088247

Pérez-Harguindeguy, N., Díaz, S., Garnier, E., Lavorel, S., Poorter, H., Jaureguiberry, P., Bret-Harte, M.S., Cornwell, W.K., Craine, J.M., Gurvich, D.E., Urcelay, C., Veneklaas, E.J., Reich, P.B., Poorter, L., Wright, I.J., Ray, P., Enrico, L., Pausas, J.G., de Vos, A.C., Buchmann, N., Funes, G., Quétier, F., Hodgson, J.G., Thompson, K., Morgan, H.D., ter Steege, H., Sack, L., Blonder, B., Poschlod, P., Vaieretti, M.V., Conti, G., Staver, A.C., Aquino, S., Cornelissen, J.H.C., 2013. New handbook for standardised measurement of plant functional traits worldwide. Aust. J. Bot. 61, 167–234.

Raffl, C., Mallaun, M., Mayer, R., Erschbamer, B., 2006. Vegetation Succession Pattern and Diversity Changes in a Glacier Valley, Central Alps, Austria. Arct. Antarct. Alp. Res. 38, 421–428. 10.1657/1523-0430(2006)38[421:VSPADC]2.0.CO;2

Ricotta, C., de Bello, F., Moretti, M., Caccianiga, M., Cerabolini, B.E.L., Pavoine, S., 2016. Measuring the functional redundancy of biological communities: a quantitative guide. Methods Ecol. Evol. 7, 1386–1395. 10.1111/2041-210X.12604

Scherrer, D., Guisan, A., 2019. Ecological indicator values reveal missing predictors of species distributions. Sci. Rep. 9, 1–8. 10.1038/s41598-019-39133-1

Scherrer, D., Lüthi, R., Bugmann, H., Burnand, J., Wohlgemuth, T., Rudow, A., 2024. Impacts of climate warming, pollution, and management on the vegetation composition of Central European beech forests. Ecol. Indic. 160, 111888. 10.1016/j.ecolind.2024.111888

Schleuter, D., Daufresne, M., Massol, F., Argillier, C., 2010. A user’s guide to functional diversity indices. Ecol. Monogr. 80, 469–484. 10.1890/08-2225.1

Shipley, B., Belluau, M., Kühn, I., Soudzilovskaia, N.A., Bahn, M., Penuelas, J., Kattge, J., Sack, L., Cavender-Bares, J., Ozinga, W.A., Blonder, B., Bodegom, P.M. van, Manning, P., Hickler, T., Sosinski, E., Pillar, V.D.P., Onipchenko, V., Poschlod, P., 2017. Predicting habitat affinities of plant species using commonly measured functional traits. J. Veg. Sci. 28, 1082–1095. 10.1111/jvs.12554

Tu, B.N., Khelidj, N., Cerretti, P., de Vere, N., Ferrari, A., Paone, F., Polidori, C., Schmid, J., Sommaggio, D., Losapio, G., 2024. Glacier retreat triggers changes in biodiversity and plant–pollinator interaction diversity. Alp. Bot. 1–12. 10.1007/s00035-024-00309-9

Venables, W.N., Ripley, B.D., 2002. MASS: modern applied statistics with S, Fourth. ed. Springer. 10.1198/tech.2003.s33

Villéger, S., Mason, N.W.H., Mouillot, D., 2008. New multidimensionale functional diversity indices for a multifaceted framework in functional ecology. Ecology 89, 2290–2301. 10.1890/07-1206.1

Walker, L.R., Wardle, D. a., Bardgett, R.D., Clarkson, B.D., 2010. The use of chronosequences in studies of ecological succession and soil development. J. Ecol. 98, 725–736. 10.1111/j.1365-2745.2010.01664.x

Wenskus, F., Hecht, C., Hering, D., Januschke, K., Rieland, G., Rumm, A., Scholz, M., Weber, A., Horchler, P., 2025. Effects of floodplain decoupling on taxonomic and functional diversity of terrestrial floodplain organisms. Ecol. Indic. 170, 113106. 10.1016/j.ecolind.2025.113106

Wright, I.J., Reich, P.B., Cornelissen, J.H.C., Falster, D.S., Groom, P.K., Hikosaka, K., Lee, W., Lusk, C.H., Niinemets, Ü., Oleksyn, J., Osada, N., Poorter, H., Warton, D.I., Westoby, M., 2005. Modulation of leaf economic traits and trait relationships by climate. Glob. Ecol. Biogeogr. 14, 411–421. 10.1111/j.1466-822x.2005.00172.x

